# SARS-CoV-2 BA.3.2.2 is more evasive of neutralization by sera from young children

**DOI:** 10.64898/2026.06.01.728533

**Authors:** Madeline Wu, Hsiang Hong, Yicheng Guo, Kristin Daniel, Ryan Hisner, Marc C. Johnson, Aubree Gordon, David D. Ho, Ian A. Mellis

**Affiliations:** Aaron Diamond AIDS Research Center, Columbia University Vagelos College of Physicians and Surgeons, New York, NY, USA; Department of Microbiology, National Taiwan University College of Medicine, Taipei, Taiwan; Department of Pathology and Cell Biology, Columbia University Vagelos College of Physicians and Surgeons, New York, NY, USA; Division of Computational Biology, Department of Integrative Biomedical Sciences, Institute of Infectious Diseases and Molecular Medicine, University of Cape Town, Cape Town, South Africa; Department of Molecular Microbiology and Immunology, School of Medicine, University of Missouri, Columbia, MO, USA; Department of Epidemiology, University of Michigan, Ann Arbor, MI, USA; Division of Infectious Diseases, Department of Medicine, Columbia University Vagelos College of Physicians and Surgeons, New York, NY, USA; Department of Microbiology and Immunology, Columbia University Vagelos College of Physicians and Surgeons, New York, NY, USA; Pandemic Research Alliance unit at the Wu Center for Pandemic Research, Columbia University Vagelos College of Physicians and Surgeons, New York, NY, USA

## Abstract

Dominant SARS-CoV-2 variants have most prominently displayed greater evasion of serum neutralizing antibodies than predecessor strains. BA.3.2, a descendant of Omicron BA.3, carrying 43 additional spike mutations, emerged in 2024, and over the last several months its subvariant BA.3.2.2 has slowly increased in prevalence globally. BA.3.2.2 continues to circulate at lower frequency than the genetically and antigenically distant dominant JN.1 subvariants NB.1.8.1 and XFG. However, concerningly, epidemiologic analyses have suggested that a larger proportion of COVID-19 cases in children are caused by BA.3.2.2 compared to adults, raising the possibility that susceptibility to BA.3.2.2 differs across age groups. Since immune imprinting shapes variant-specific anti-SARS-CoV-2 antibody profiles and children born after 2021 primarily were first exposed to Omicron subvariants, we hypothesized that young children may have lower circulating neutralizing antibody titers against BA.3.2.2 than adults. Using pseudovirus neutralization assays, we measured titers against BA.3.2.2 and other SARS-CoV-2 variants in serum or plasma samples from a total of 36 adults (≥18 years old), school-age children (3-10 years old), and infants/toddlers (6-28 months old) in the US. We found that both cohorts of children had lower geometric mean titers against BA.3.2.2 than adults, even though all tested age groups had similar titers against dominant strains NB.1.8.1 and XFG. Together, these findings suggest that susceptibility to emerging SARS-CoV-2 variants may diverge across age groups, perhaps as a result of their different exposure histories. Furthermore, these results highlight the importance of SARS-CoV-2 surveillance and the monitoring of immunity against viral variants across age ranges.

## Main Text

Since the start of the COVID-19 pandemic, successive SARS-CoV-2 variants have driven repeated waves of immune escape, necessitating periodic reevaluation of vaccine composition and other public health measures^1,2^. Occasionally, saltation or recombination events have generated highly divergent spikes capable of reshaping viral spread, most recently exemplified by the emergence of the now dominant JN.1 sublineages^3^.

BA.3.2, a highly divergent descendant of the early-Omicron BA.3 lineage carrying 43 additional spike mutations, emerged in 2024 in South Africa. Over the last several months, its subvariant BA.3.2.2 has slowly increased in global prevalence (**Figure A**)^4,5^. Since March 1, 2026, limited available surveillance data suggest that BA.3.2.2 may have grown to represent a majority or plurality of reported SARS-CoV-2 sequences in some locations (e.g., 75/88 sequences from Spain, 53/73 from Italy, 22/37 from Malaysia). Unusually, epidemiologic analyses of samples submitted for sequencing suggest that BA.3.2.2 may disproportionately infect young children compared to adults **(Figure B)**, raising the possibility that susceptibility to this lineage differs across age groups with distinct SARS-CoV-2 exposure histories^4^. Such substantial enrichment in pediatric cases was not observed for XFG and NB.1.8.1 (**Figure S1A**). Considered across variants, the fraction BA.3.2.2 sequences derived from children is higher than the fractions for XFG and NB.1.8.1 (**Figure S1B**). We note, however, that any differences in asymptomatic infection frequency between variants may lead to reporting biases.

**Figure.**
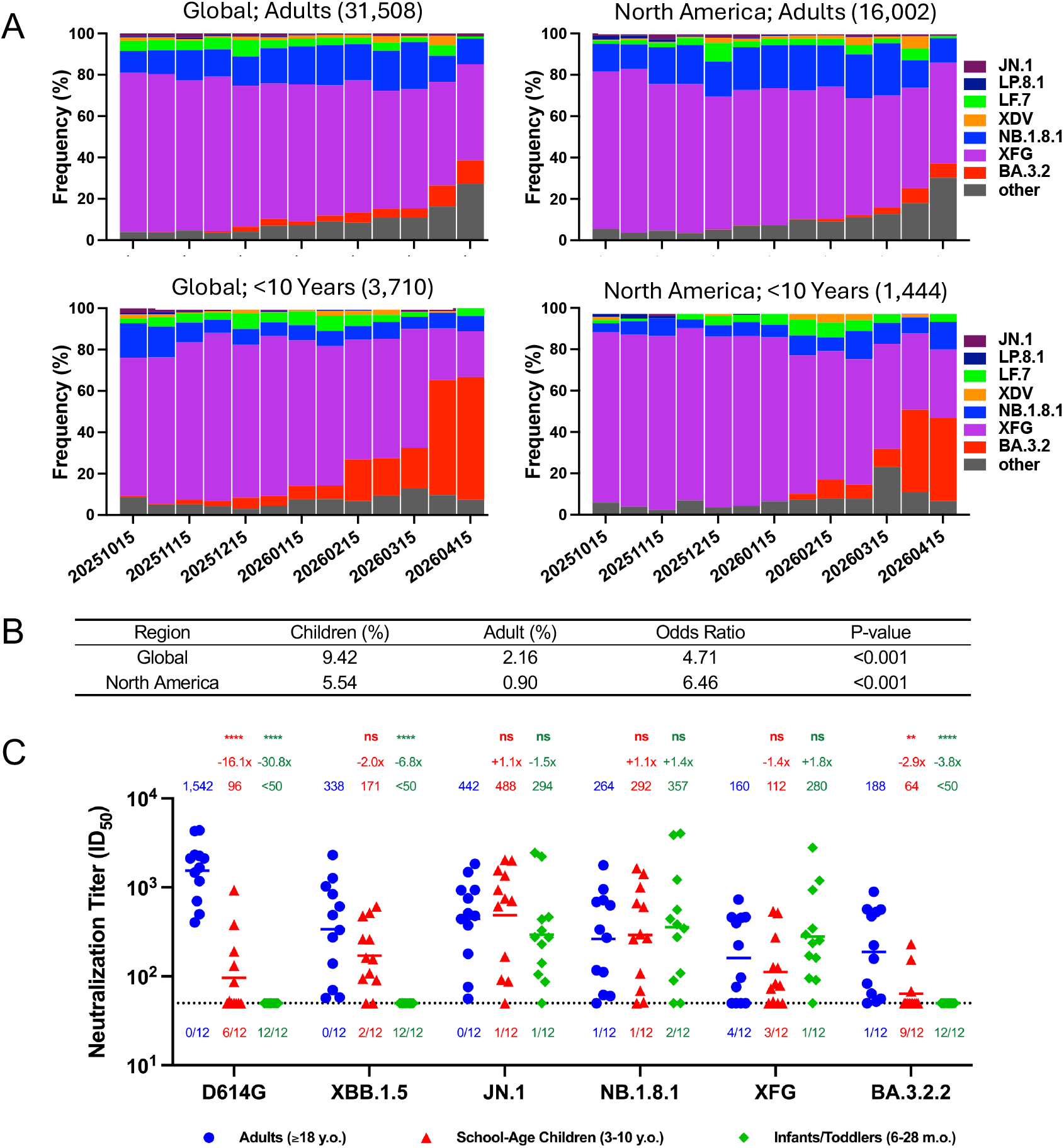
Frequency of and neutralizing antibody titers against SARS-CoV-2 BA.3.2.2 in young children. a. Relative frequencies of SARS-CoV-2 variants from October 15, 2025, to April 15, 2026, stratified by age group and region. Top panels show adults (≥18 years) and bottom panels show children (<10 years). Left panels represent global sequences and right panels represent North American sequences. Numbers in parentheses indicate sequences with available age information. Most BA.3.2 sublineage sequences are the BA.3.2.2 subvariant. b. Statistical analysis of BA.3.2 sublineage frequencies globally and in North America. Statistical significance was assessed using a two-sided two-proportion z-test comparing the proportion of BA.3.2 sequences between children aged <10 years and adults aged ≥18 years. Global frequencies were 337/3690 (9.13%) versus 678/31,432 (2.16%), respectively; North American frequencies were 66/1401 (4.71%) versus 140/15,821 (0.88%). c. Serum or plasma pseudovirus neutralization titers (ID_50_) against SARS-CoV-2 BA.3.2.2 and other variants in adults (≥18 years old), school-age children (3-10 years old), and infants/toddlers (6-28 months old). Dashed line indicates limit of detection (LOD) = 50. 12 samples per group were tested for neutralization, and number of samples below LOD listed under dashed line. Geometric mean titers (GMT) indicated above points, and fold differences compared to adult titers are shown above GMT. Asterisks corresponding to Mann Whitney U test p-values shown above fold differences. ****: p<0.0001; ***: p<0.001; **: p<0.01; *: p<0.05; ns: not significant.

Since immune imprinting shapes variant-specific anti-SARS-CoV-2 antibody profiles^6^ and children born after 2021 primarily were first exposed to Omicron subvariants whereas older children and adults were first exposed to earlier strains, we hypothesized that young children may have lower neutralizing antibody titers against BA.3.2.2 than adults. To characterize titers across age ranges, we evaluated pseudovirus neutralization with serum or plasma samples from 12-person cohorts of adults (≥18 years), school-age children (3-10 years), and infants/toddlers (6-28 months), tested against D614G, XBB.1.5, JN.1, NB.1.8.1, XFG, and BA.3.2.2 (**Figure C)**. We chose 3 years old as the age threshold between pediatric cohorts to separate those born before the introduction of monovalent Omicron XBB.1.5-directed vaccines (3-10 years), who may have had an ancestral SARS-CoV-2 exposure through vaccination or infection, and those born after (6-28 months), who likely did not.

There were no significant age-specific differences in geometric mean titers (GMT) against XFG or NB.1.8.1 across the 3 cohorts. Adults had neutralizing activity against BA.3.2.2 (GMT = 188, and only one sample below the limit of detection [LOD]), that was comparable to their titers against the dominant variant XFG (GMT=160). However, neutralization of BA.3.2.2 was markedly lower than XFG in school-age children (BA.3.2.2 GMT=64; 9/12 below LOD) and undetectable in infants/toddlers. The 3 children with detectable BA.3.2.2-neutralizing titers had detectable D614G-directed titers, suggesting that they may have had more adult-like exposure histories (**Figure S2**). Notably, the tested plasma from the infants/toddlers demonstrated neutralization only against JN.1, NB.1.8.1, and XFG, suggesting a more lineage-specific neutralization profile rather than uniformly weak SARS-CoV-2 antibody responses. Re-grouping the cohorts into those who were more likely to have any pre-Omicron exposures (i.e., born before mid-2021) and those who were not, revealed a similar trend in variant-specific titers (**Figure S3**).

These findings suggest that susceptibility to emerging SARS-CoV-2 variants may diverge across age groups with different exposure histories. Adults in the US and many other countries have accumulated broader immunity through repeated infection and vaccination across antigenically distinct lineages, starting with the ancestral strain, whereas younger children and infants possess narrower exposure histories largely shaped by recent variants. Continued surveillance of SARS-CoV-2 variants should consider age-stratified differences in immunity, to anticipate or explain disproportionate burden of infections in some populations.

## Notes

### Competing Interest Statement

D.D.H. co-founded TaiMed Biologics and RenBio, and he serves as a consultant for Brii Biosciences and is a board director at Vicarious Surgical. A.G. served as a member of the scientific advisory board for Janssen Pharmaceuticals and has consulted and serves on a scientific advisory board for Sanofi Pasteur.

### Funding

- National Institutes of Health: subcontract no. 0258-A700-4609 under federal contract no. 75N93021C00014, subcontract GR0010139-PO024016 under federal contract no. 75N93021C00016, K08AI196255
- Columbia University: UR014016
- Heart of Racing

