## Supplementary Appendix for "SARS-CoV-2 BA.3.2.2 is more evasive of neutralization by sera from young children"

### Methods

#### Clinical Cohorts

De-identified residual whole blood samples from children who had presented for lead testing by the clinical laboratories of NewYork-Presbyterian/Columbia University Irving Medical Center were procured through the Columbia University Center for Advanced Laboratory Medicine (CALM) in accordance with the protocol AAAO2000 approved by the Columbia University IRB. CALM, as the laboratory's honest broker, selected and de-identified residual samples from children at least 6 months old and up to 10 years old who did not have a documented concurrent positive test result for SARS-CoV-2. Upon receipt, samples were centrifuged, plasma was aliquoted, and samples were screened for SARS-CoV-2 viremia by RT-PCR and excluded if positive. Then, samples were screened for the presence any anti-SARS-CoV-2 spike-binding antibodies using ELISA and included only if any spike-binding antibodies were detected. ELISA screening enabled more direct comparison against the adult cohort, who are known to have prior COVID-19 vaccination and/or SARS-CoV-2 infection histories. Participants in the school-age children cohort were 58% female, 42% male, with an average of 4.7 years. Participants in the infants/toddlers cohort were 41.7% female, 58.3% male, with an average age of 1.7 years.

Adult serum samples drawn in late 2025 before a vaccine booster dose were collected through the VIVA study at the University of Michigan and through the "COVID-19 Persistence and Immunology Cohort (C-PIC)" study at Columbia University. Specimens were obtained following formal participant informed consent, adhering to the protocols approved by the IRBs of University of Michigan Medical School (protocol HUM00232359) and Columbia University (protocol AAAS9722). Participants in the adult cohort were 75% female, 25% male, with an average age of 34.3 years.

Details are summarized in **Table S1**. All serum and plasma samples were heat inactivated at 56°C for 30 minutes before use.

**Cell Lines**

Vero-E6 (CRL-1586) cells and HEK293T (CRL-3216) cells were obtained from ATCC and cultured at 37°C with 5% $CO_2$ in Dulbecco's Modified Eagle Medium (DMEM) + 10% fetal bovine serum (FBS) + 1% penicillin-streptomycin. Vero-E6 cells are derived from African-green monkey kidneys. HEK293T cells are of human female origin.

**Plasmid Generation**

As previously described, in plasmids for pseudoviruses, mutations were made using the QuickChange II XL and QuickChange Multi site-directed mutagenesis kits (Agilent). All constructs were verified using whole plasmid sequencing prior to use.

**Pseudovirus production**

VSV-based SARS-CoV-2 pseudoviruses, in which the native VSV glycoprotein was replaced by SARS-CoV-2 spike and its variants, were produced as previously described. Briefly, plasmids containing the appropriate spike were transfected into HEK293T cells with PEI. After 24 hours, VSV-G pseuotyped ΔG-luciferase (G*ΔG-luciferase, Kerafast) was added, and then washed with medium three times before being cultured in fresh medium for another 24 hours. Anti-VSVG (anti-I1) antibody was added to deplete non-pseudotyped viruses. Pseudoviruses were then harvested, centrifuged, and then aliquoted and stored at -80°C.

**Pseudovirus neutralization assays**

Each SARS-CoV-2 pseudovirus was titrated to standardize viral infectious dose before use in neutralization assays. Seven serial dilutions of heat-inactivated sera or plasma were added in 96-well plates, starting at 1:50 dilution. Next, pseudoviruses were added and incubated at 37 °C for 1 hour. In each plate, wells containing only pseudoviruses were included as controls. $4 \times 10^4$ Vero-E6 cells were then added per well and incubated at 37 °C for 16 hours. Cells were lysed and luminescence was determined by the Luciferase Assay System (Promega E4550) and Tecan Infinite® 200 PRO using i-control™ software v.3.9.1.0, in accordance with the manufacturer's instructions. The serum dilution that inhibits 50% of virus entry ($ID_{50}$) was calculated using nonlinear five-parameter dose-response curve fitting using GraphPad Prism v.10.3.1.

**Quantification and statistical analysis**

The serum dilution that inhibits 50% of virus entry ($ID_{50}$) was determined by fitting a five-parameter dose-response curve in GraphPad Prism v.11.3.1 and in R v4.3.2 using the drda v2.0.3 package.

### Author Contributions

The study was conceptualized by I.A.M. and D.D.H. Samples were collected and/or requested by I.A.M. and A.G. Experiments were conducted and data analyzed by M.W., H.H., Y.G., K.D., and I.A.M. Initial demographic analyses were conducted by R.H. and M.J. The manuscript was written by M.W., H.H., Y.G., I.A.M., and D.D.H. All contributing authors have received and endorsed the manuscript.

### Acknowledgements

We express our gratitude to Hiroshi Mohri (Columbia) for assistance with sample collection, to Jayesh G. Shah, Lawrence J. Purpura, Amanda Castillo, Meredith McNairy and Antonia Sturiza for conducting the C-PIC study (Columbia), and to Carmen Gherasim, Virginia M. Pierce, Theresa Kowalski-Dobson, Anna Buswinka, Joseph Wendzinski, Mayurika Patel, Noah Paalanen and Ethan Hall of the VIVA study team for conducting the VIVA study. We thank Erin Poptanich, Tiffany Thomas, and Eldad Hod for advice and assistance related to requesting residual pediatric blood samples. We also thank all who share data on GISAID. This study was supported by funding from the NIH SARS-CoV-2 Assessment of Viral Evolution (SAVE) Program (subcontract no. 0258-A700-4609 under federal contract no. 75N93021C00014 to D.D.H and subcontract GR0010139-PO024016 under federal contract no. 75N93021C00016 to A.G.), K08AI196255 salary support to I.A.M., internal startup funding UR014016 from Columbia University to Y.G., and Heart of Racing to M.C.J.

### Declaration of Interests

D.D.H. co-founded TaiMed Biologics and RenBio, and he serves as a consultant for Brii Biosciences and is a board director at Vicarious Surgical. A.G. served as a member of the scientific advisory board for Janssen Pharmaceuticals and has consulted and serves on a scientific advisory board for Sanofi Pasteur.

### Supplementary Figures and Table

A

| region | strain | Children (%) | Adult (%) | Odds Ratio | p value |
|---|---|---|---|---|---|
| Global | XFG | 67.18 (2,496/3,715) | 68.94 (21,723/31,512) | 0.922 | 0.029651 |
| Global | NB.1.8.1 | 9.07 (337/3,715) | 14.89 (4,695/31,512) | 0.57 | 7.91E-22 |
| North America | XFG | 70.94 (1,011/1,425) | 65.68 (10,430/15,879) | 1.28 | 5.79E-05 |
| North America | NB.1.8.1 | 7.65 (109/1,425) | 18.54 (2,944/15,879) | 0.36 | 5.06E-25 |

Statistical analysis of XFG and Nb.1.8.1 sublineage frequencies globally and in North America. Statistical significance was assessed using a two-sided two-proportion z-test comparing the proportion of BA.3.2 sequences between children aged <10 years and adults aged ≥18 years.

B

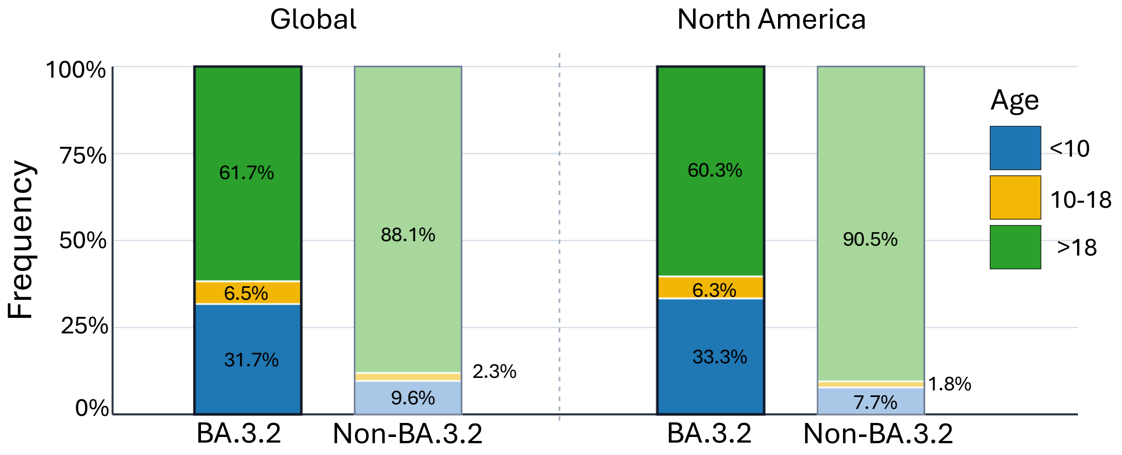

**Figure S1: Variant frequencies across age ranges.**

a. Statistical analysis of XFG and NB.1.8.1 sublineage frequencies globally and in North America. Statistical significance was assessed using a two-sided two-proportion z-test comparing the proportion of BA.3.2 sequences between children aged <10 years and adults aged ≥18 years.

b. Only sequences with an annotated patient age range were considered. Stacked bars show the age composition of BA.3.2 and non-BA.3.2 SARS-CoV-2 sequences globally and in North America, from October 2, 2025 to May 1, 2026. Samples were stratified into three age groups: <10 years, 10-18 years, and ≥18 years. BA.3.2 bars are shown with saturated colors, while non-BA.3.2 bars are shown with lighter tints. Bar heights represent the percentage contribution of each age group within the indicated sequence group. Data from GISAID.

| Sample | Age (Years) | Neutralization Titers ($ID_{50}$) | | | | | |
|---|---|---|---|---|---|---|---|
| | | D614G | XBB.1.5 | JN.1 | NB.1.8.1 | XFG | BA.3.2.2 |
| CALM 24 | 0.75 | 50 | 50 | 105 | 109 | 170 | 50 |
| CALM 21 | 1.00 | 50 | 50 | 230 | 444 | 253 | 50 |
| CALM 20 | 1.25 | 50 | 50 | 141 | 1221 | 95 | 50 |
| CALM 15 | 1.33 | 50 | 50 | 438 | 384 | 295 | 50 |
| CALM 16 | 1.50 | 50 | 50 | 2222 | 4029 | 2800 | 50 |
| CALM 23 | 1.83 | 50 | 50 | 275 | 276 | 346 | 50 |
| CALM 19 | 1.92 | 50 | 50 | 276 | 90 | 91 | 50 |
| CALM 4 | 2.00 | 50 | 50 | 50 | 50 | 50 | 50 |
| CALM 11 | 2.00 | 50 | 50 | 2443 | 3887 | 936 | 50 |
| CALM 2 | 2.08 | 50 | 50 | 329 | 346 | 237 | 50 |
| CALM 8 | 2.17 | 50 | 50 | 465 | 560 | 1190 | 50 |
| CALM 12 | 2.17 | 50 | 50 | 87 | 50 | 162 | 50 |
| CALM 25 | 3 | 50 | 611 | 88 | 51 | 125 | 50 |
| CALM 27 | 3 | 50 | 478 | 2049 | 1644 | 518 | 50 |
| CALM 29 | 3 | 50 | 50 | 1360 | 1422 | 52 | 50 |
| CALM 31 | 3 | 50 | 94 | 167 | 108 | 79 | 50 |
| CALM 33 | 4 | 191 | 91 | 752 | 270 | 274 | 68 |
| CALM 39 | 4 | 50 | 262 | 1560 | 604 | 126 | 50 |
| CALM 46 | 5 | 87 | 163 | 2013 | 1000 | 539 | 231 |
| CALM 49 | 5 | 50 | 109 | 92 | 69 | 50 | 50 |
| CALM 36 | 6 | 132 | 261 | 943 | 263 | 50 | 50 |
| CALM 37 | 6 | 382 | 50 | 50 | 50 | 50 | 50 |
| CALM 38 | 7 | 53 | 521 | 616 | 300 | 83 | 50 |
| CALM 40 | 7 | 931 | 156 | 711 | 667 | 73 | 154 |
| CUMC 3 | 22 | 701 | 70 | 179 | 62 | 50 | 83 |
| CUMC 8 | 23 | 1465 | 332 | 442 | 117 | 76 | 158 |
| CUMC 16 | 23 | 1181 | 274 | 481 | 271 | 50 | 56 |
| CUMC 18 | 23 | 1666 | 487 | 751 | 685 | 453 | 223 |
| CUMC 1 | 26 | 2261 | 839 | 929 | 952 | 398 | 563 |
| MICH 6 | 26 | 501 | 57 | 76 | 60 | 50 | 52 |
| MICH 2 | 30 | 4291 | 1272 | 510 | 336 | 223 | 566 |
| MICH 3 | 31 | 4371 | 2305 | 1837 | 1771 | 463 | 892 |
| MICH 5 | 33 | 403 | 58 | 56 | 50 | 50 | 50 |
| MICH 1 | 48 | 2320 | 612 | 934 | 720 | 732 | 528 |
| MICH 7 | 52 | 2121 | 1028 | 1486 | 630 | 462 | 475 |
| MICH 4 | 75 | 2117 | 139 | 375 | 112 | 97 | 64 |

**Figure S2: Neutralizing antibody titers of all participants sorted by age.** Serum or plasma pseudovirus neutralizating titers (ID50) against SARS-CoV-2 BA.3.2.2 and other variants in adults (≥18 years old), school-age children (3-10 years old), and infants/toddlers (6-28 months old). Color scale denote neutralizing titers.

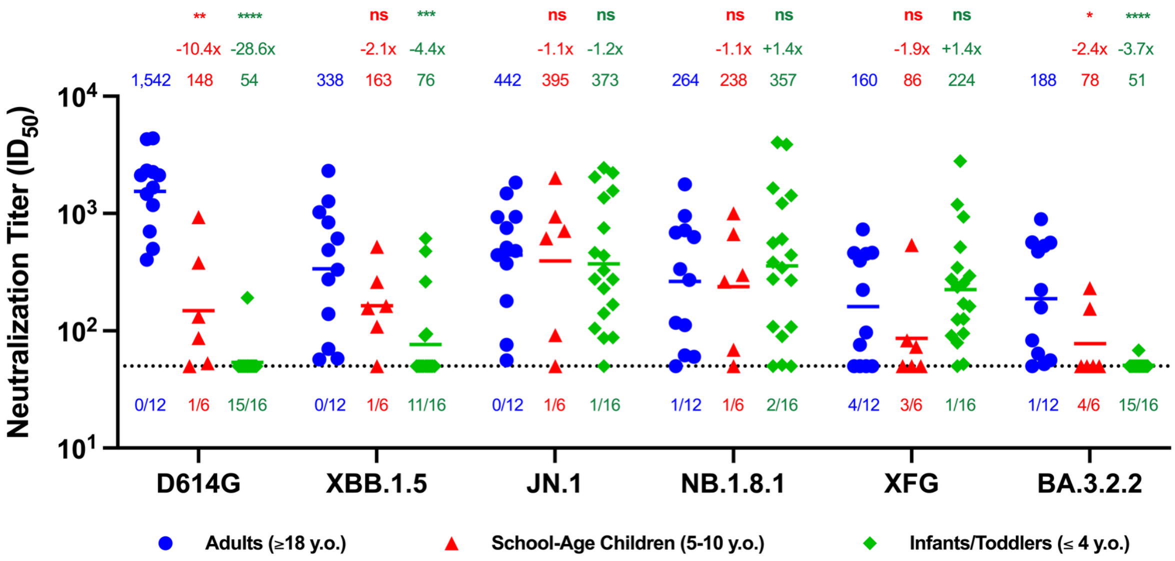

**Figure S3: Neutralizing antibody titers among participants born before or after the emergence of SARS-CoV-2 Omicron.** Serum or plasma pseudovirus neutralizating titers ($ID_{50}$) against SARS-CoV-2 BA.3.2.2 and other variants in adults (≥18 years old), children aged 5 to 10 years old, and children age 6 months to 4 years old. Only those with BA.3.2.2 $ID_{50}$ > 50 included. Dashed line indicates limit of detection (LOD) = 50. Number of samples below LOD listed under dashed line. Geometric mean titers (GMT) indicated above points, and fold differences compared to adult titers are shown above GMT. Asterisks corresponding to Mann Whitney U test p-values shown above fold differences. ****: $p<0.0001$; ***: $p<0.001$; **: $p<0.01$; *: $p<0.05$; ns: not significant.

| | All participants | | Adults (18+ y.o.) | | School-age Children (3-10 y.o.) | | Infants/Toddlers (6-28 m.o.) | |
|---|---|---|---|---|---|---|---|---|
| | No. or Mean | % or (range) | No. or Mean | % or (range) | No. or Mean | % or (range) | No. or Mean | % or (range) |
| **Total** | 36 | - | 12 | - | 12 | - | 12 | - |
| **Female** | 21 | 58.3% | 9 | 75.0% | 7 | 58% | 5 | 41.7% |
| **Male** | 15 | 41.7% | 3 | 25.0% | 5 | 42% | 7 | 58.3% |
| **Age (y.o.)** | 13.6 | (0, 75) | 34.3 | (22, 75) | 4.7 | (3, 7) | 1.7 | (9m, 26m) |

**Table S1: Cohort summary.** y.o.: years old; m.o.: months old.
